# Modular and state-relevant connectivity in high-frequency resting-state BOLD fMRI data: An independent component analysis

**DOI:** 10.1101/2020.07.22.212720

**Authors:** Thomas DeRamus, Ashkan Faghiri, Armin Iraji, Oktay Agcaoglu, Victor Vergara, Zening Fu, Rogers Silva, Harshvardhan Gazula, Julia Stephen, Tony W. Wilson, Yu-Ping Wang, Vince Calhoun

**Affiliations:** Tri-institutional Center for Translational Research in Neuroimaging and Data Science (TReNDS), Georgia State University, Georgia Institute of Technology, Emory University, Atlanta, Georgia, USA; Department of Electrical and Computer Engineering, Georgia Tech, Georgia, USA; The Mind Research Network, Albuquerque, NM, USA; Department of Neurological Sciences, University of Nebraska Medical Center, Omaha, Nebraska; Center for Magnetoencephalography, University of Nebraska Medical Center, Omaha, Nebraska; Department of Biomedical Engineering, Tulane University, New Orleans, LA, USA; Department of Psychology, Georgia State University, Georgia, USA

**Keywords:** Blood-oxygenation-level-dependence frequency spectrum analysis, spatially constrained independent component analysis, resting-state functional magnetic resonance imaging

## Abstract

Resting-state fMRI (rs-fMRI) data are typically filtered at different frequency bins between 0.008∼0.2 Hz (varies across the literature) prior to analysis to mitigate nuisance variables (e.g., drift, motion, cardiac, and respiratory) and maximize the sensitivity to neuronal-mediated BOLD signal. However, multiple lines of evidence suggest meaningful BOLD signal may also be parsed at higher frequencies. To test this notion, a functional network connectivity (FNC) analysis based on a spatially informed independent component analysis (ICA) was performed at seven different bandpass frequency bins to examine FNC matrices across spectra. Further, eyes open (EO) vs. eyes closed (EC) resting-state acquisitions from the same participants were compared across frequency bins to examine if EO vs. EC FNC matrices and randomness estimations of FNC matrices are distinguishable at different frequencies.

Results show that FNCs in higher-frequency bins display modular FNC similar to the lowest frequency bin, while r-to-z FNC and FNC-based measures indicating matrix non-randomness were highest in the 0.31-0.46 Hz range relative to all frequency bins above and below this range. As such, the FNC within this range appears to be the most temporally correlated, but the mechanisms facilitating this coherence require further analyses. Compared to EO, EC displayed greater FNC (involved in visual, cognitive control, somatomotor, and auditory domains) and randomness values at lower frequency bins, but this phenomenon flipped (EO > EC) at frequency bins greater than 0.46 Hz, particularly within visual regions.

While the effect sizes range from small to large specific to frequency range and resting state (EO vs. EC), with little influence from common artifacts. These differences indicate that unique information can be derived from FNC between BOLD signals at different frequencies relative to a given restingstate acquisition and support the hypothesis meaningful BOLD signal is present at higher frequency ranges.

## Introduction

Resting-state functional network connectivity (FNC) derived from brain networks informed by spatial independent component analysis (sICA), has become a ubiquitous approach for assessing brain function and probing diagnostic biomarkers. The signal of interest in most resting-state functional connectivity (FC) or FNC studies is typically temporally filtered up-to the ∼0.2 Hz range (Biswal, DeYoe, & Hyde, 1996), with the purpose of mitigating the contribution of nuisance signal in BOLD timeseries, including respiration (0.3-1.0 Hz), cardiovascular activity (≥1 Hz), scanner drift (≤ 0.01 Hz), and motion (varies) (Biswal et al., 1996; Cordes et al., 2001; Niazy, Xie, Miller, Beckmann, & Smith, 2011; Satterthwaite et al., 2013). However, there is recent evidence indicating BOLD signal in higher frequencies up to the limit allowed by the sampling rate (1/[2*Repetition Time]) might also be of neuronal origin (Chen & Glover, 2015).

On the other hand, (Chen, Jahanian, & Glover, 2017) and others have noted many studies examining FNC patterns across broader frequency ranges did not correctly pre-process their datasets. It is suggested that high-frequency findings are primarily driven by sequential nuisance regression of artifact (Chen et al., 2017; Lindquist, Geuter, Wager, & Caffo, 2019). This issue calls into question results from previous analyses of high-frequency data and their relevance for extracting diagnostically useful information. In this regard, FNC patterns within these higher frequency ranges have not been adequately studied.

FC/FNC structure generally reveals “modular patterns,” or a grid-like organization with discrete and overlapping temporal and/or topological properties (Ferrarini et al., 2009; Newman, 2006; Sporns, 2003; Yu et al., 2011). It has been posited that alterations within such systems reflect changes in functional brain states in humans due to clinical or developmental alterations in functional brain organization (Burger & Schuler, 2012; Chen et al., 2017; Yuan, Wang, Zang, & Liu, 2014). Similar arguments have been made for functional network alterations in clinical populations in higher frequencies within the BOLD spectra. However, few of these studies have specifically examined modularity across a range of frequencies, nor have they studied what factors may influence these functional network connections. Such differences in FC/FNC across the frequency spectra may provide useful information in clinical populations (Morgan, Rogers, & Abou-Khalil, 2015; Sours et al., 2015; Wang et al., 2015).

Studies where participants serve as their own controls while the same filtering approaches are applied across different resting-state paradigms would provide a clear window into the behavior of FNCs derived from higher frequency BOLD signal. Previous studies have demonstrated significant alterations in FNC between eyes open (EO) vs. eyes closed (EC) resting state (Agcaoglu, Wilson, Wang, Stephen, & Calhoun, 2019; Liu, Dong, Zuo, Wang, & Zang, 2013; Patriat et al., 2013; Wang et al., 2015; Yuan et al., 2014; Zou et al., 2009) using a standard low or bandpass filter (typically between 0.008∼0.2). As such, we investigate FNC between eyes open (EO) vs. eyes closed (EC) resting state in various frequency bands.

For this purpose, the authors utilized a spatially constrained method of independent component analysis (ICA) (Du & Fan, 2013; Li & Adali, 2010a, 2010b; Ma et al., 2011) to extract intrinsic connectivity networks (ICNs) across sub-sections of the frequency spectra. Five-minute time series of resting-state functional magnetic resonance imaging (rs-fMRI), with participants both lying in the scanner with EC and EO conditions (counter-balanced across individuals), are utilized for this objective. The primary goal of this research is to evaluate the presence of non-random FNC patterns among high-frequency BOLD signals. This is achieved by analysis of FNC matrices within each frequency bin using the matrix randomness proposed by Vergara and colleagues (Vergara & Calhoun, 2016; Vergara, Yu, & Calhoun, 2018a, 2018b). As a secondary objective, the authors explore differences in FNC modularity between EO and EC rs-fMRI within and across each frequency bin.

## Methods

### Participants

Participants include 174 youth with no reported clinical diagnoses from the greater Albuquerque, New Mexico (92) and Omaha, Nebraska (82) areas as part of the NSF funded collaborative developmental chronnecto-genomics consortium (Dev-CoG; http://devcog.mrn.org/), between the Mind Research Network (MRN), University of Nebraska Medical Center (UNMC), and Tulane University. All parents signed informed consent forms, and youth signed assent forms approved by each participating university’s institutional review board before participating in the study. The appropriate institutional review board for each study site approved all procedures. Of the participants, 89 reported as male, 84 reported as female, and one did not report sex or gender. No significant differences were identified regarding age and Full-Scale Intelligence Quotient (FSIQ) between participants who reported as male or female. However, comparisons between acquisition sites identified significantly higher FSIQs in participants from UNMC compared to MRN. There were no other significant differences across age or gender composition between MRN and UNMC. The comparisons are summarized in Table 1.

**Table 1.** Demographic comparisons of participants

|  | N | Range | Difference (*W*) | Means(*t*) | Test Statistic | *p*-value | *CI*95% | *Eff* size |
| --- | --- | --- | --- | --- | --- | --- | --- | --- |
| **F vs M** | 89/84 |  |  |  |  |  |  |  |
| Age |  | 9.2-15.1 vs 9.1-15.5 | -0.1 |  | *W* = 3607 | *p* = 0.691 | -0.7 : 0.4 | *r* = -0.03 |
| FSIQ |  | 72-140 vs 68-148 |  | 108.605 vs 112.138 | *t*<sub>(165.429)</sub> = -1.557 | *p* = 0.121 | -8.014 : 0.948 | *g* = -0.238 |
| **MRN vs UNMC** | 92/82 |  |  |  |  |  |  |  |
| Age |  | 9.1-15.5 vs 9.4-15.5 | -0.2 |  | *W* = 3520 | *p* = 0.531 | -0.8 : 0.4 | *r* = -0.048 |
| F vs M | 46/46 vs 38/43 | | | | $\chi^2_{(1,174)} = 0.0639452$ | *p* = 0.8 | | $\phi = 0.031$ |
| *FSIQ |  | 72-139 vs 68-148 | -0.883 | 108.272 vs 112.813 | *t*<sub>(165.205)</sub> = -2.002 | *p* = 0.046 | -9.0179 : -0.062 | *g* = -0.307 |

Note: \* Indicates statistical significance at or below a 0.05 threshold.
*W* represents the rank-difference statistic from a Wilcoxon rank-sum test, quantified using the difference in ranks and an *r* effect size
*t* represents the difference in means derived from a Welch's two or paired sample *t*-test, effect size is quantified using Hedge's *g*
$\chi^2$ Represents the goodness-of-fit from a Pearson chi-squared test. Effect size is displayed using $\phi$ .
*V* Represents the Wilcoxon signed-rank test pseudo median, effect size is quantified using a paired *r*.
Positive *g* & *r* values indicate greater values in females or MRN, while negative values indicate greater values males or UNMC.

### Imaging data collection

Imaging data collected at MRN utilized a 3T Siemens TIM Trio scanner, while a 3T Siemens Skyra scanner was used at the UNMC site. One single-band-reference image *(SBref)*, followed by a total of 650 volumes rs-fMRI echo-planar imaging BOLD data were collected for both EC and EO for each participant. Rs-fMRI scans were acquired using a standard gradientecho echo-planar imaging paradigm with a repetition time (TR) of 0.46 s, echo time (TE) = 29 ms, flip angle (FA) = 44°, and a slice thickness of 3 mm with no gap. The acquisition parameter for the MRN site used a field of view (FOV) of 246 × 246 mm (82 × 82 matrix), and 56 sequential axial slices, while the UNMC site utilized a FOV of 268 × 268 mm (82 × 82 matrix), and 48 sequential axial slices. The order of the EO and EC resting-state sessions were counter-balanced across participants at each site.

### Data Quality Analyses

Analyses were conducted on images from both EC and EO conditions for each participant to assess potential differences due to data quality. The median value of the mean time series image divided by the standard deviation of the time series, the standard deviation of outlier voxels by image (calculated using *AFNI’s 3dToutcount)*, head motion (quantified using the Euclidean norm [ENORM] using *AFNI’s 3dvolreg)*, and inherent smoothness *(FWHM;* computed with *AFNI’s 3dFWHMx)* were compared across gender, site, and resting-state paradigm. Participants identifying as female displayed significantly greater median *tSNR* and motion/*ENORM* during the EC condition, while participants reporting as male exhibited greater motion/*ENORM* during the EO condition. Median *tSNR* and inherent smoothness were significantly greater in the MRN site compared to the UNMC site, while motion and outlier voxel counts did not significantly differ between sites. Finally, EC and EO only differed on inherent smoothness and no other calculated metrics. The results of each statistical test are reported in Supplementary Table 1 and Supplementary Figure 1.

**Figure 1.**
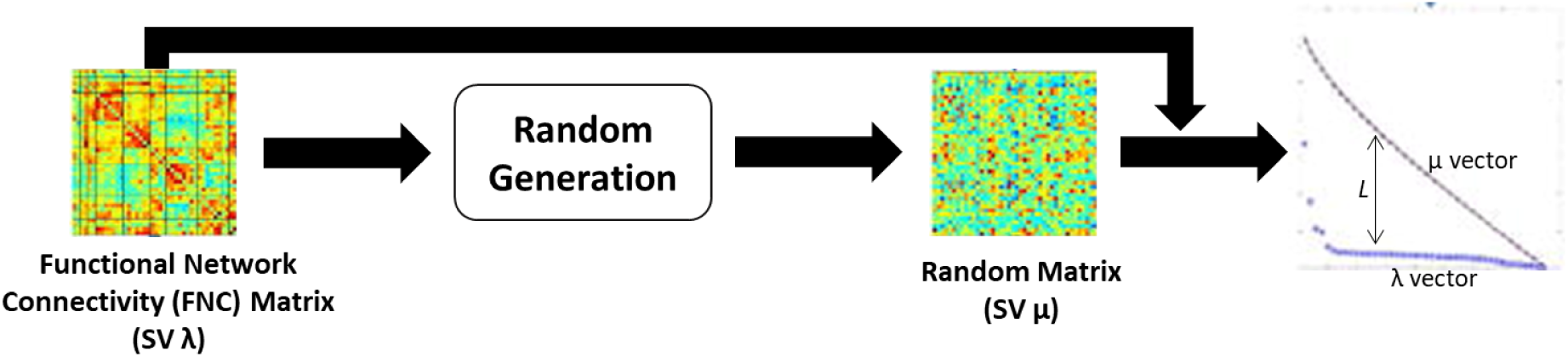
(Left)A functionaI network connectivity (FNC) with 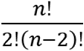 pairwise component Combinations scaled to singular values (SV) λ. (Middle) A randomly generated, gaussian-based connectivity matrix, scaled to SV μ. (Right)The *L* distance between the plotted SVs for the randomly generated (upper μ plot), and FNC SV λ plot. A greater value for *L* indicates less randomness.

### Preprocessing

Images were pre-processed using a hybrid pipeline of *FSL 5*.*0*.*5* (Jenkinson, Beckmann, Behrens, Woolrich, & Smith, 2012; Smith et al., 2004), *AFNI* (Cox, 1996) and *ANTs* (Brian B. Avants, Tustison, & Song, 2009). First, images were distortion-corrected using FSL’s *topup* using AP-PA phase encoding. Second, *ANTs’ antsMultivariateTemplateConstruction2*.*sh* (Brian B Avants, Tustison, Wu, Cook, & Gee, 2011; Brian B Avants, Tustison, Song, et al., 2011; Sanchez, Richards, & Almli, 2012) was used to create a custom pediatric template from the distortion-corrected single-band reference *(SBRef)* images. Third, the first 10 of the 650 images in the distortion-corrected timeseries for each participant were removed to facilitate magnetization equilibrium relative to the first scans. Fourth, the distortion-corrected timeseries were then despiked using *AFNI’s 3dDespike* -NEW flag for multi-band data to increase registration accuracy (Caballero-Gaudes & Reynolds, 2017; Jo et al., 2013). Fifth, the data were realigned to the first image in the timeseries. Finally, the first image in each participant’s timeseries was non-linearly registered using ANT’s greedy *SyN* diffeomorphic registration (B B Avants, Epstein, Grossman, & Gee, 2008; Klein et al., 2009) to a 3mm isometric custom template constructed from all participant’s single-band reference *(SBRef)* images. Transformations were computed from the study-specific template to an MNI-space template constructed from warps of averaged timeseries data transformed directly to an isometric 3mm MNI space (via *SPM8;* (Friston et al., 1994), and participant-to-template and template-to-MNI transformations were performed on the first image in each timeseries for all participants. No coregistration to anatomical images was performed in this workflow. Following normalization, *AFNI’s 3dTproject* was utilized to apply a regression model to simultaneously perform: 1) linear detrending, 2) second- and third-order Legendre polynomials to remove nuisance variables across the timeseries, 3) sine and cosine functions modeling frequencies outside the evenly spaced bins between 0.01-1.76 Hz (mimicking a passband filter using regression) and smoothing the residuals with a 6mm *FWHM* kernel. The residuals of BOLD timeseries from each set of 640 images (for each resting-state paradigm for each participant) were variance normalized (timeseries are linearly detrended and converted to z-scores for each voxel) prior to ICA.

### Independent Component Analysis

The network templates used for the current study were computed from previous work by (Agcaoglu et al., 2019), who performed ICA on the full frequency spectrum of the data (no filters were applied prior to ICA) using the *GIFT toolbox* (http://trendscenter.org/software/gift/) in *MATLAB*. First, EC and EO scans were pooled across participants, and principal component analyses (PCAs) were employed for each participant timeseries to reduce the dimensionality across 646 (as opposed to 640 for the current study) time points to 200 maximally variable directions. Second, the principal components for each participant for EC and EO were concatenated across the time dimension, and a group PCA was applied to further reduce the dimensionality to 150 maximally variable directions (Erhardt et al., 2011). Third, 150 independent components were estimated from the group PCA matrix with the Infomax algorithm (Bell & Sejnowski, 1995), repeated 20 times in *ICASSO* (Himberg & Hyvarinen, 2003) (http://www.cis.hut.fi/projects/ica/icasso) with the most central runs selected from the resulting 20 runs to ensure the stability of the estimated components (Ma et al., 2011).

The workflow utilizes a spatially constrained multivariate objective optimization ICA with reference (MOO-ICAR) to estimate sources using component maps as spatial priors (Du & Fan, 2013; Du et al., 2016). For this study, the group-level independent component analysis (gICA; (Calhoun, Adali, Pearlson, & Pekar, 2001); (Calhoun et al., 2001) components from Agcaoglu et al., 2019 (see supplementary material for an HTML-based component viewer) were used to extract independent component maps and their time courses from EC and EO for each frequency bin (Agcaoglu et al., 2019). Fifty components were determined to be intrinsic connectivity networks components were determined to be domain-associated intrincsic (functional) connectivity networks (DA-ICNs) using visual inspection and atlas labels from the “whereami” function in *AFNI* at the peak coordinates for each component. These include: 3 auditory (AUD), 2 cerebellar (CB), 12 cognitive control (CC), 4 default mode (DM), 4 subcortical (SC), 8 somatomotor (SM), and 17 visual (VS) components. DA-ICN matrices from each pairwise combination of DA-ICNs totaled to 1225 possible combinations, with 20% consisting of within ICN (e.g. one DM component to another DM component), and 80% constituting between ICN (e.g. DM to SC components) correlations.

Functions from the *GIFT toolbox* (http://trendscenter.org/software/gift/) in *MATLAB* were deployed within in-house workflows distributed across the TReNDS high-performance computing (HPC) environment (for specifications, see: https://help.rs.gsu.edu/display/PD/Advanced+Research+Computing+Technology+and+Innovation+Core+%28ARCTIC%29+resources) using a *SLURM* (Yoo, Jette, & Grondona, 2003) scheduler and resource manager. Software frameworks, versions, and builds by the study are listed in Supplementary Table 2.

### Randomness Estimations

As described in (Vergara & Calhoun, 2016; Vergara et al., 2018a, 2018b), the randomness of FNC matrices can be quantified using the mathematical framework first set forth by Girko (1985) which is now known as the circle law (Girko, 1985; Marčenko & Pastur, 1967; Trotter, 1984). Unlike graphtheory based measures (Rubinov & Sporns, 2010), the value associated with each edge in a random matrix is not weighted or binarized but is assumed to follow a Gaussian distribution with properties like the Fisher r-to-z transformed correlation. The randomness test fails (depending on hypothesis testing and statistical significance) if matrix edge values do not follow a Gaussian random distribution. The proposed randomness test utilizes the characteristics of the FNC matrix using matrix singular value-spectrums. For each participant’s FNC matrix, a singular value decomposition (SVD) is computed. Assessing matrix randomness is achieved by comparing the FNC singular values (SVs) of each participant against the SVs of random matrices, where a participant’s FNC SVs are compared against SVs of randomly generated matrices M with Gaussian matrix elements. The distance between the singular FNC vector and the randomly generated vector (*L*) computed using *Equation 1*

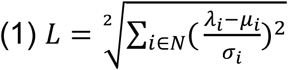

where *N* is the total number of functional networks which make up the matrix of interest (for each participant), *i*. is a member of *N* matrix, *μ*_*i*_ represents the mean and *σ*_*i*_ the standard deviation of the FNC matrix of interest, and *λ*_*i*_ is the vector with SVs for the (participant) FNC matrix of interest. Figure 1 adapted from (Vergara et al., 2018a) displays a brief illustration of the relationship between variables in the singular value spectra. The computations were performed using in-house *MATLAB* workflow distributed across the TReNDS HPC environment as described previously in the ICA section.

Once the *L* values are recorded for each participant across each resting-state paradigm and frequency bin, a single linear mixed effect (LME) model is performed on all *L* values using the *lmer* function in the *lme4* (Bates, Mächler, Bolker, & Walker, 2015) *R* package (R Core team, 2015; Team, 2019). A permutation-based LME model with age, gender, FSIQ, EC vs. EO order, data acquisition site (MRN vs. UNMC), resting-state paradigm (EC vs. EO), frequency bin, motion, the interaction of frequency and motion, and the interaction between frequency bin and resting-state paradigm using the *permanova*.*lmer* function in the *lmerTest* and *predictmeans R* packages (Bates et al., 2015; Kuznetsova, Brockhoff, & Christensen, 2017; Luo, Ganesh, & Koolaard, 2020) were utilized to perform the LME. The *permanova*.*lmer* produces 5000 permutations of *L* values to generate a *p*-distribution for the data for significance testing when the assumption of normally distributed residuals is not met. As the frequency bin is the factor of interest, and the effect of resting-state paradigm is shared within participants, pairwise comparisons of the residuals between frequencies were split across (with the effects of all other variable regressed out) resting-state paradigm and *Holm* corrected across all comparisons. The *R* version and package builds are also listed in Supplementary Table 2.

### Linear Mixed-Effect model for FNC matrices

Several LME models were fit to each pairwise *r-to-z* FNC set of values within the FNC matrix using the same *lmer* function in the *lme4* (Bates, Mächler, Bolker, & Walker, 2015) package in *R*, producing a total of 1225 individual analyses. This approach was selected to minimize the number of multiple comparison corrections across all frequency bins (7*1225 for *FNC)*, for individual cell-wise regressions or pairwise contrasts. Like the previous model used to test randomness, the FNC based *LME* model included participants as a random effect, and age, gender, FSIQ, acquisition site, restingstate paradigm, resting-state paradigm order, frequency bin (with 0.01-0.15 Hz as the reference), motion calculated using ENORM, and the interaction between frequency bin and motion as fixed effects. *P*-distributions for each FNC pairing were estimated using the *permanova*.*lmer* function within the *predicmeans* (Luo et al., 2020) package, with 5000 permuted FNC values per pairing. The effect of frequency on the FNC pairings between networks serves as the variable of interest and is false discovery rate (FDR) corrected at the level of the *p*-value for the omnibus *F*-value for each frequency bin across all FNC pairs. Interactions with motion and frequency bin (and other variables of no interest) are reported at factor-wise FDR corrected *p*-values to identify potential confounds and their influence on the model. The FDR corrected *p*-values by factor results for these variables are presented in the supplemental material.

Pairwise comparisons across the residuals of FNC pairs across frequency bin with nuisance variables removed were assessed using a Wilcoxon signed-rank test (R Core team, 2015; Team, 2019; Wilcoxon, 1946). Significant interactions between frequency bin and motion are examined using post-hoc paired sample Ptests of slopes using the *emmeans* (Lenth, 2019) and *interactions* (Long, 2019) packages in *R* using the *emtrends* function within the *emmeans* library.

As with the models for the *L* variable computed in equation 1, pairwise comparisons between frequency residuals were split across resting-state paradigm and *Holm* corrected (Holm, 1979; Lenth, 2019; Torchiano, 2016) across all comparisons as the frequency bin is the effect of interest and the effect of resting-state paradigm is shared within participants. *R* version and package builds are described in Supplementary Table 2.

## Results

### Randomness

The LME model identified statistically significant differences by frequency bin (*F*_(6, 2163.5)_ = 254.40, *p*_perm_ = 0.0002,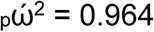), acquisition site (*F*_(1, 162.1)_ = 8.79, *p*_perm_ = 0.0028, 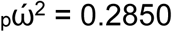), restingstate paradigm (*F*_(1, 2163.5)_ = 13.73, *p*_perm_ = 0.0056, 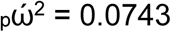), the interaction of resting-state paradigm by frequency (*F*_(6, 2163.5)_ = 4.7695, *p*_perm_ = 0.0004, 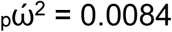) and the interaction of frequency bin and motion (*F*_(6, 2163.5)_ = 4.84, *p*_perm_ = 0.0002, 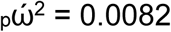), but not motion (*F*_(1, 338.80)_ = 0.0018, *p*_perm_ = 1, 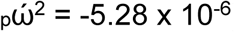), age (*F*_(1, 162.1)_ = 0.2677, *p*_perm_ = 0.6106, 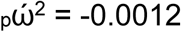, gender (*F*_(1, 162.1)_ = 0.0041, *p*_perm_ = 1, 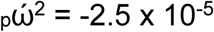), FSIQ (*F*_(1, 162.1)_ = 0.0182, *p*_perm_ = 1, 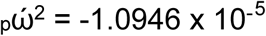), or resting-state paradigm order (*F*_(1, 2169.5)_ = 0.0098, *p*_perm_ = 0.9314, 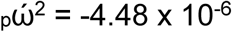). The *conditional R*^*2*^ of 0.4697, and a *marginal R*^*2*^ of 0.3680, suggests a relatively short distance between the observed values and those fitted by the LME model (Johnson, 2014; Nakagawa & Schielzeth, 2013).

#### Cellwise Comparisons of Frequency Bins

Pairwise comparisons between frequency bins (averaged across resting-state paradigm) were not significantly different between one another even at an uncorrected threshold, despite the statistically significant effect of frequency bin in the *LME* model.

#### EC vs. EO Across Frequency Bins

Like the frequency bin comparisons, contrasts of EC vs. EO (averaged across all frequency bins) did not return as statistically significant *(p* = 0.7568, *r* = 0.0034).

#### Resting-State Paradigm by Frequency Bin Interaction

When compared across the same frequency bins, significantly increased *L* distance (indicating less matrix randomness) was found in the EC matrices for the 0.61 -0.76 and 0.91 -1.07 Hz bins (*p* = 0.0409, *g* = 0.0046, *p* = 0.006, *g* = 0.0064), but no other pairings. Across frequency bins, FC was noticeably less random in EC compared to EO across most of the contrasts 0.31-0.46 Hz, the exceptions of which occurring at the lower frequency bins (0.01-0.31 Hz), and when EO from the 0.31-0.46 Hz frequency bin was compared to any EC not within the 0.31-0.46 Hz bin. These results are summarized in Figure 2a

**Figure 2.**
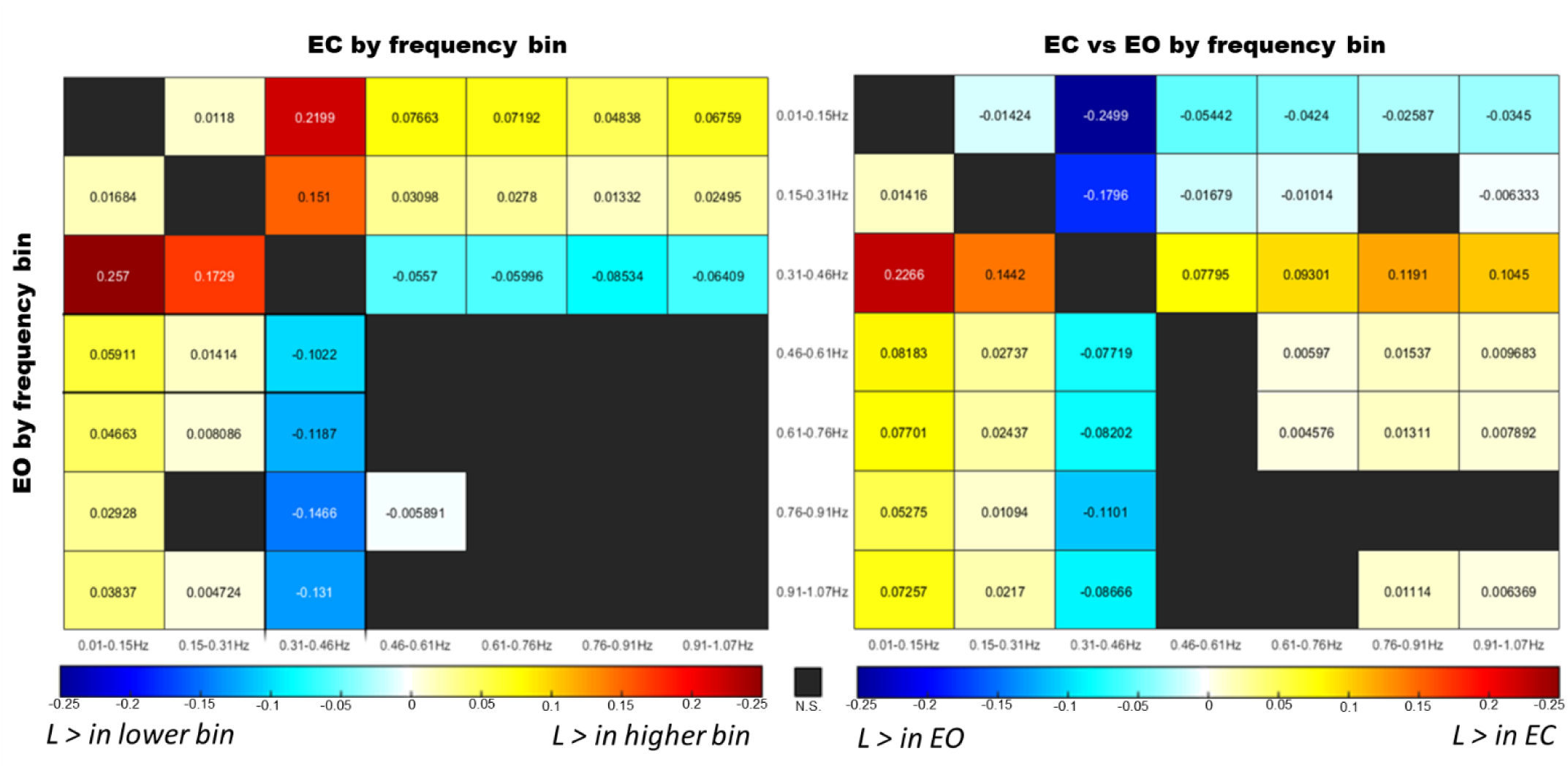
Comparisons of between-task randomness *(L)* residuals across frequency bands. Metric displayed is the Hedge’s *g* for each contrast. (Left) Comparisons *L* residuals between resting-state paradigm with EC in the upper triangular and EO on the Iower triangular. Cool Colors indicate a higher *L* value in the Iower frequency bin, while warm colors indicate a higher *L* value in the higher frequency bin. (Right) EC vs EO Comparisons with EC as the reference across the seven frequency bins. Warm colors indicate greater *L* values in the EC condition, while cool colors indicate greater *L* values in EO.

When comparing within resting-state paradigm, few differences were noticeable when frequency bins above 0.46 Hz were compared to one another, the exception occured between 0.46-0.61 vs. 0.911.07 Hz in EO (with the former having a greater *L*/less randomness). For frequency bins below 0.46 Hz, most contrasts favored a greater *L* in the higher frequency bin relative to the lower bins. The exception to this was at the 0.46-0.61 Hz bin, which displayed a significantly increased *L*/less randomness across comparisons with all other bins. These results are displayed in Figure 2b.

#### Pairwise Artifact Comparisons

Motion measured using the ENORM did not significantly influence the LME model used for *L*. However, significantly greater randomness in motion-by-frequency-bin slopes were found in the 0.01-0.15 Hz (*p*_Holm_ = 4.032 × 10^−4^, *g* = 0.0110) and 0.15-0.31 Hz (*p*_Holm_ = 0.0140, *g* = 0.0079) relative to the 0.31-0.46 Hz bin. These results are depicted in Supplementary Figure 2. No significant differences in data acquisition sites were noted when contrasts were averaged across frequency bin and restingstate paradigm.

### FNC comparisons

The marginal (fixed-effect only) *R*^*2*^ for the models ranged from 0.0282-0.5435 (mean = 0.1916), while the conditional (random and fixed-effects) *R*^*2*^ for the models ranged from 0.1578-0.6517 (mean = 0.3645), suggesting wide variability in model variance by FC pairs (Jaeger, 2017; Johnson, 2014; Nakagawa & Schielzeth, 2013). The linear mixed effect models found all the 1225 FNC pairs significantly influenced (following *FDR* correction) by the frequency bin through which the data was filtered prior to gig-ICA. Partial ω^2^ values for frequency bin were highly variable, ranging from 0.0003-0.9910, with 1090 (89%) meeting the criteria for a large effect for frequency bin when variance from all other independent variables are removed. For contrasts between EC vs. EO (averaged across all frequency bins), roughly 712 (∼58%) of the models were significantly influenced by resting-state paradigm, with 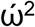 values ranging 0.0028-0.9761, with 267 (∼23%) meeting the criteria for a large effect size. When examining the interactions between frequency bin by resting-state paradigm, 774 (∼63%) of the 1225 models were significantly influenced by the relationship between resting-state paradigm and frequency bin, with 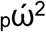 values ranging from 0.0006-0.4867, and 27 (∼2%) displaying large effect sizes 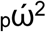. Figure 3 displays both *R*^2^ and 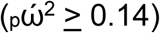 values for bin and resting-state paradigm for all models across FNC pairs Statistically significant effects were identified for the site of data acquisition, with 965 pairings (∼79%) having significant effects ranging from 0.0075-0.9919 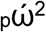, 745 of which met the criteria for large effect sizes. ENORM motion significantly influenced the LME models of 555 FC pairs (∼45%), with 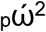 ranging from 0.0166-0.9504, 293 of which met criteria for a large effect size. Interactions between frequency bin and motion were identified for 836 (∼68%) FC pairs, with effect sizes ranging from 0.0004-0.3314, but only 5 of which met criteria for large 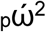. These findings are discussed further in the pairwise comparisons section below.

**Figure 3.**
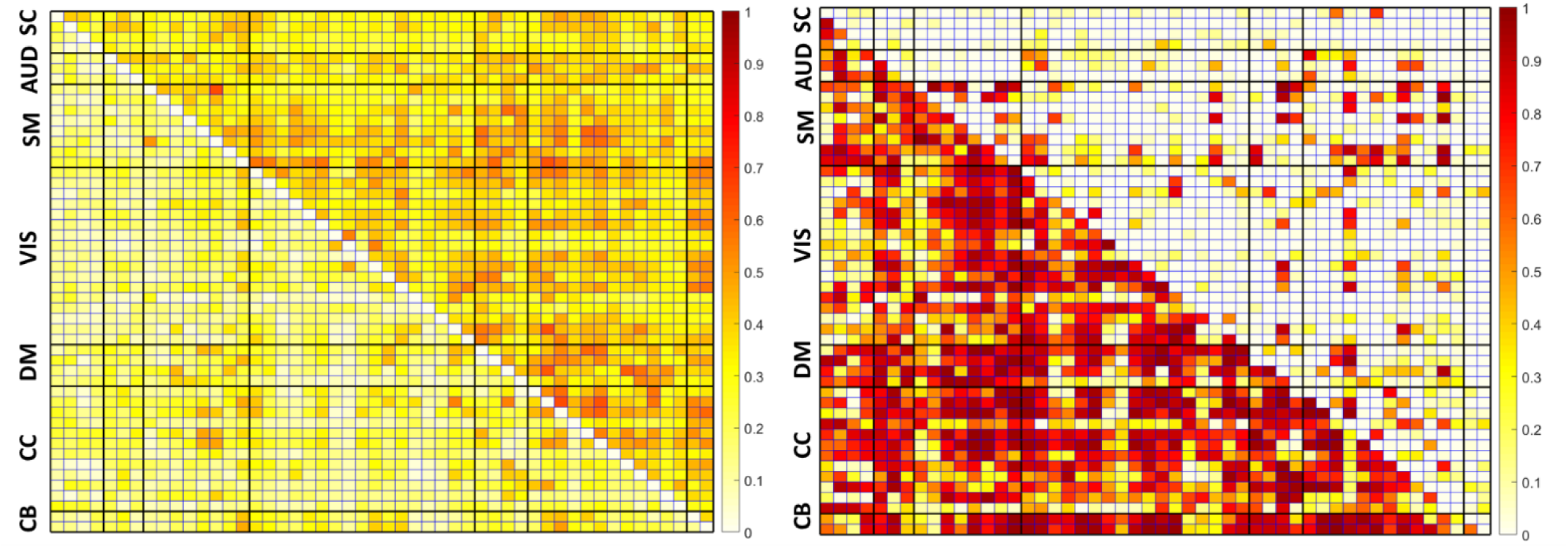
FNC matrices displaying (left)*R*^2^ marginal (fixed-effects only Iower triangular) and *R*^*2*^ conditional (fixed and random effects. Upper triangular) indicative of model fit (right) indicating 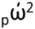 for Statistically significant effects for frequency band (lower triangular) and resting-state paradigm (upper triangular).

#### Pairwise Comparisons

The average r-to-z transformation for EC and EO values are displayed on the top row in Figure 4. None of the subsequent pairwise analyses across frequency bins (averaged across resting-state paradigm) or resting-state paradigm (averaged across frequency bins) displayed statistically significant differences following pairwise EC vs. EO contrasts for resting-state paradigm, or *Holm* corrected comparisons across frequency bins averaged across resting-state paradigms (21 contrasts). While pairwise comparisons of the main effects of resting-state paradigm or frequency bin did not appear significant when averaged, multiple contrasts of resting-state paradigm across frequency bin (91 comparisons per FC pair) were statistically significant following a *Holm* correction. The bottom row of Figure 4 depicts the *Pearson r* effect sizes of all within-frequency contrasts of EC vs. EO.

**Figure 4.**
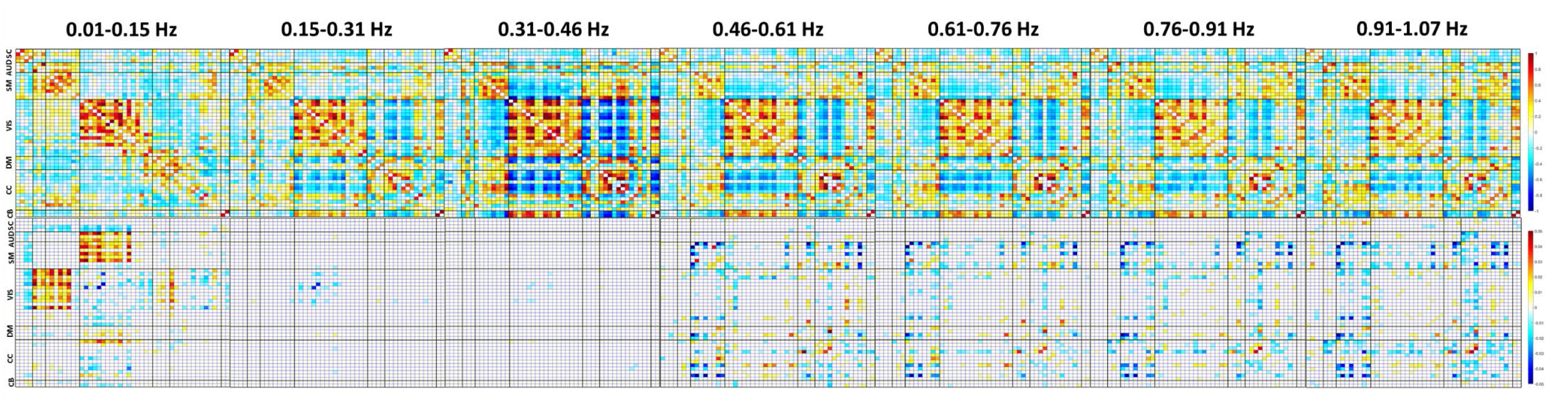
(Top) The color indicates the average value of the positive (red) or negative (blue) correlation represented with the r-to-z transform. The lower triangular matrix is averaged across EC, while the upper is averaged across EO. (Bottom) FNC matrices of *r* effect sizes for FDR corrected LME models and Holm corrected pairwise tests between EC and EO within frequency bins, *+r* (red) indicates the value is greater in EC condition, while *-r* (blue) indicates greater values in the EO condition. Note the differences in scaling between the average FC values and effect sizes.

For contrasts across frequency bins for EC, within-network connections were more likely to be larger r-to-z transformed FC measures in lower frequency bins for SC, SM, VS, and DMN regions, with the opposite displayed in AUD and CC regions. For EO, within-network connections were more likely to have increased r-to-z values in lower frequencies in SC, AUD, SM, and DMN pairs, while larger r-to-z values FC in were observed in VIS and CC connections within higher frequency bins. Between-network connections in EC were more likely to have larger r-to-z values in lower frequencies in SC, AUD, SM, DMN, and CC regions, and larger r-to-z values in higher frequencies in VIS and CB connections. For EO, lower frequencies were more likely to display increases in r-to-z values across SC, AUD, and CC regions, and larger r-to-z values in higher frequency bins in SM, VS, DMN, and CB pairings. The distribution of these results quantified using difference scores in total results is displayed in Figure 5.

**Figure 5.**
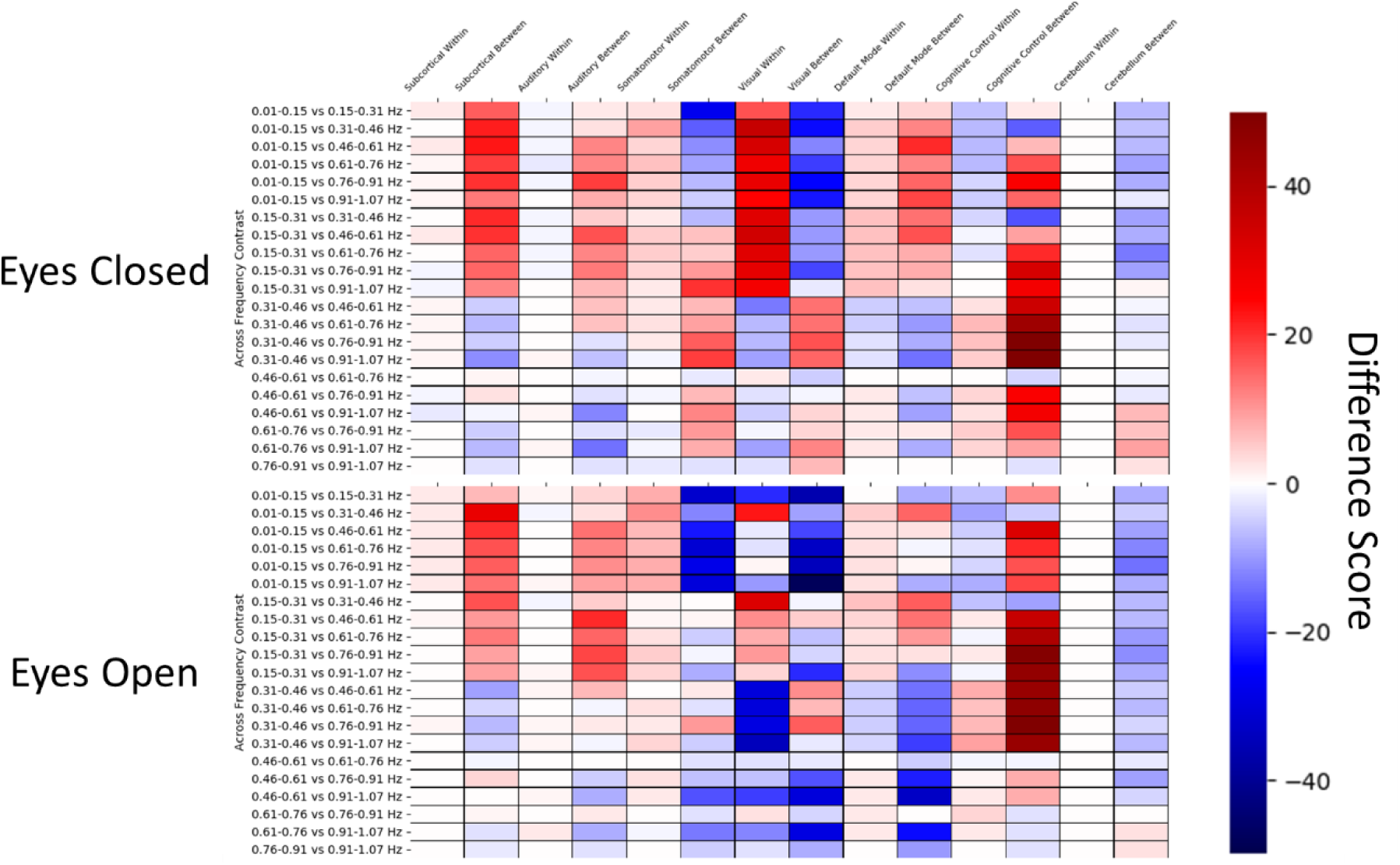
Difference scores between the total number of Connections within or between **a**network to quantify trends across frequency bins. Warm colors indicate that of the significant results, the network is more Iikely to trend towards the Iower frequency compared to the higher frequency, while cool colors indicate the total number of significant results tend to favor greater FC in the higher frequencies. The Top figure display the counts across the eyes open condition, while the bottom displays the eyes closed condition.

When examining contrasts of EC vs. EO across frequency bins, EC was more likely to be greater within SC, AUD, and CC FC pairs and between AUD and CC pairs. EO was typically found to be greater within-network SM, VIS, and DMN, and between network SC, SM, VIS, DMN, and CB connections. These results are summarized using difference scores across all contrasts and quantified by the total number of comparisons across all connections in Figure 6.

**Figure 6.**
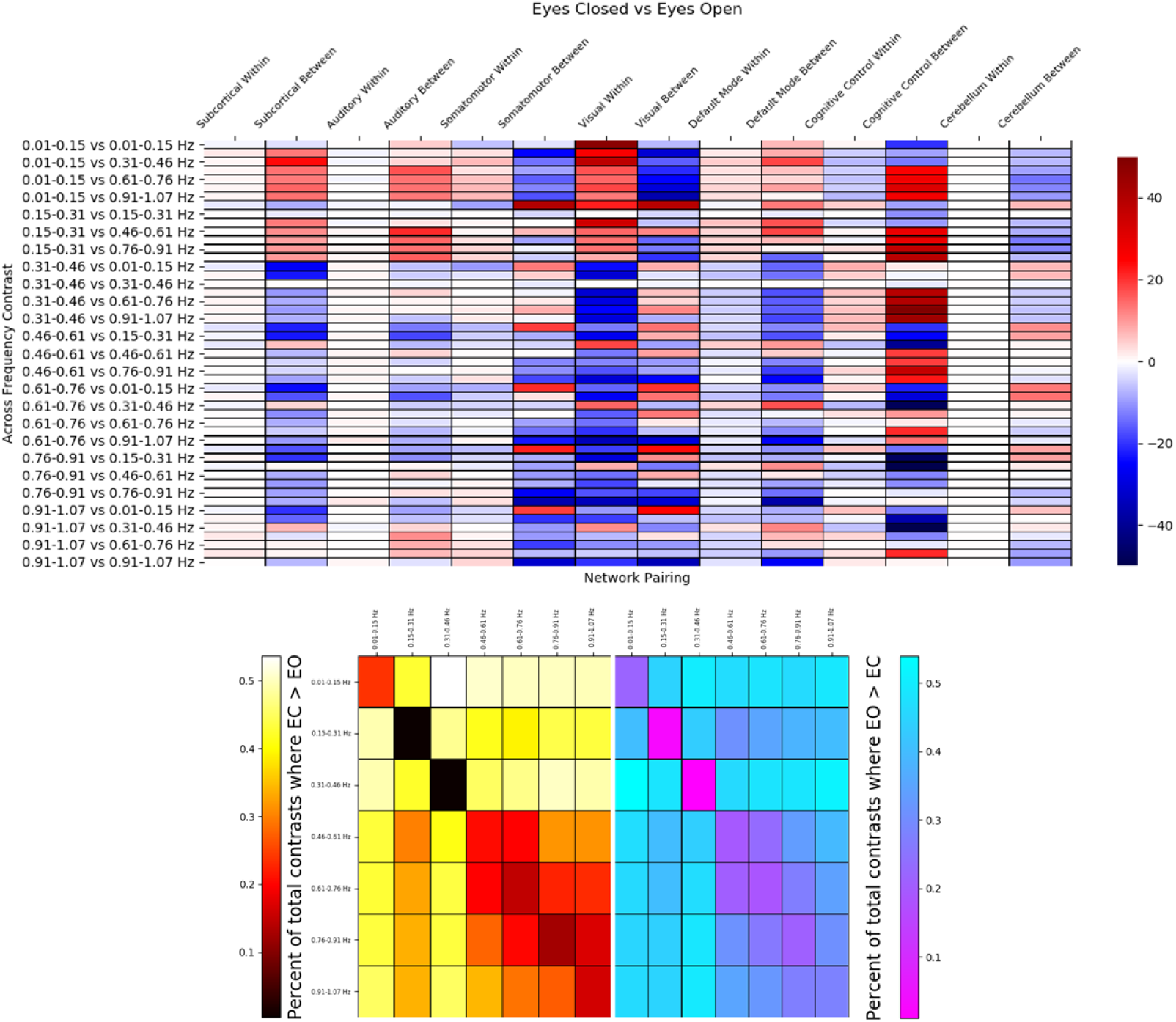
(Top) Difference scores by statisticalIy significant result counts by network. Warm colors indicate that a Contrast trends towards FC vaIues being greater in EC, while cool Colors indicate greater values in EO. (Bottom) Quantification of the percentages of significant results by the total number of Contrasts across each frequency bin comparison. The left graph with the warm colors display Contrasts where effects are greater in EC compared to EO, while the figure on the right and the cool colors indicate where EO is greater to EC. Note that the scale is in raw percentages, not fractions of the total.

The significant changes in slopes with frequency bin and motion are presented with the statistically significant slopes between frequency bin and resting-state paradigm as reference referenced in Figure 7. Motion-related slopes were more likely to be greater in lower frequencies compared to higher frequencies in all types of FNC pairs except for within AUD and CB pairings.

**Figure 7.**
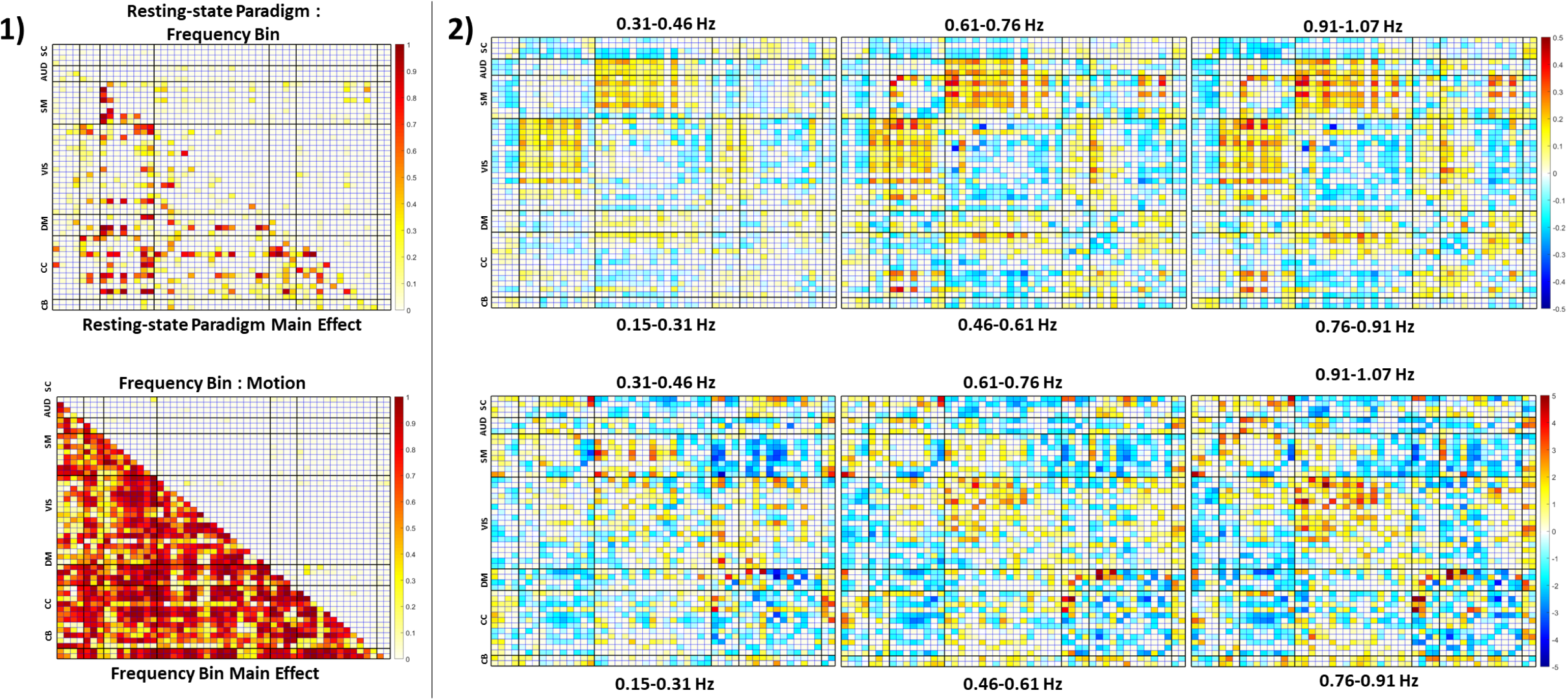
1) FNC matrices of partial ω^2^ from the linear mixed effect models. The top image displays the partial ω^2^for the main effect of resting-state paradigm (Iower trianguIar), and interaction between frequency bin and resting-state paradigm (upper triangular). The bottom figure displays the ω^2^ value for the main effect of frequency bin (Iower trianguIar) relative to the interaction between frequency bin and motion (Upper trianguIar). 2) Top: The slopes for the interactions of resting-state paradigm by each frequency bin relative to the 0.01-0.15 frequency bin. Bottom: The slopes of the interaction between frequency bin and motion each frequency bin above 0.01-0.15 Hz. The slopes are unstandardized, note the differences in scale between the frequency bin slopes (−0.5, 0.05), and frequency bin by motion interaction (−5,5).

## Discussion

The results of non-randomness of FNC matrices suggest meaningful functional connectivity information can be captured at high-frequency BOLD signals (up to the Nyquist frequency limit) higher than the low-frequency ranges commonly used in rsfMRI FNC analyses. Furthermore, the results indicate modularity is preserved at higher frequencies across brain networks and display variable patterns. The FNC non-randomness appears to be greater in lower frequencies compared to higher frequencies, peaking at 0.31-0.46 Hz before decreasing and displaying negligible differences across higher frequencies above 0.46 Hz. However, with the exception of the 0.01-0.15 Hz bin in the traditional filtering frequency, differences between EC and EO resting-state conditions are considerably less prevalent at lower frequencies compared to higher frequencies. These results suggest that the information provided within a given frequency range is dependent upon the question of interest. The interaction between resting-state paradigm and frequency bin support this claim. Should the information in distinct BOLD frequency ranges be neuronal in origin, FNC matrices and other methods could serve as useful neural correlates for estimating FC coupling strength and differences across resting-state paradigms at different frequencies.

Consistent with (Agcaoglu et al., 2019), there were significant differences in FNC between EO and EC at the lowest frequency bin (0.01-0.15 Hz). Interestingly, few differences are found in lower-to-middle frequency ranges, but were prevalent at frequencies above 0.46 Hz. VIS, CC, and SM ICNs were more likely to be influenced by resting-state condition. This finding is interesting but not surprising considering the length of research illustrating amplitude differences in visual and somatomotor brain regions specific to EC vs EO (Liang et al., 2014; Wei et al., 2018; Zou et al., 2015). However, these results are likely the only EC vs. EO findings to date in frequency bins outside of traditional lower frequencies employed in fMRI analyses. As such, it is possible the results presented here at the higher-frequencies may represent neuronally-influenced occipital and somatomotor recordings mediated by resting-state paradigm. Were this the case, nuanced analyses specific to such systems could be tailored with a combination of resting-state paradigm and filtering method.

It is worth noting several FNCs were significantly influenced by average participant motion even after correction for multiple comparisons. Most of these findings occurred in higher-frequencies relative to lower frequencies, and were more prevalent in SC, AUD, SM, VIS, and CC domains pairings. This is of particular concern in the present study as all of these regions are prone to significantly greater influence of respiration artifact and motion-related distance dependence artifact (see (Burgess et al., 2016) for a brief overview). As such, even though there appeared to be little effect of motion in isolation, careful consideration should be applied to interpretations made regarding results within these domains. Future work would benefit from combinations of ICA, nuisance spectra (Fair et al., 2020; Gratton et al., 2019) and data-driven optimization not only to isolate signal of interest, but also design objective filters to minimize artifact, as artifact and signal of interest may not be compartmentalized to the evenly-spaced frequency bins selected for this analysis. Were such approaches to be utilized, they would need to be robust to acquisition parameters such as the Nyquist frequency limits set by the TR, and signal-and-artifact frequency overlap. Work by (Vergara & Calhoun, 2020) have demonstrated data-driven filters for targeting signals-of-interest in focused frequency ranges, and the application of such approaches will be critical but for evaluating high-frequency BOLD studies.

Our study, for the first time to the authors’ knowledge, to examined differences in high-frequency BOLD FNC analyses within a spatially constrained ICA framework. While it has been lightly examined in (Chen et al., 2017), the focus of that study was on high-frequency contamination from sequential nuisance regression, an approach that was eschewed in within the current study. The results presented in this work provide a starting point for the development and application of tools to assess and tune BOLD-based resting-state paradigm and FNC analyses, with the potential to expand into temporal dynamics once specific trends have been isolated and replicated across datasets. Further, the “bin-based” approach common within the high-frequency literature may be sub-optimal and implementing data-driven methods for filtering may expand on the results presented here.

### Limitations

As with any fMRI study, the effects of motion may significantly contribute to the measured BOLD signals, particularly long vs. short range FNC values. Furthermore, recent work has identified that short TR multi-band sequences have respiration artifacts that have alias with motion estimates, and without information specific to respiration (e.g., waist bellows), notch filtering is recommended to mitigate respiration (Fair et al., 2020; Gratton et al., 2019). For the current study, however, motion measures likely mixed with respiration were fitted as a single regressor, both to avoid the loss of frequency-specific information associated with notch filtering, and because respiration information was not collected in the dataset. The authors suggest that a follow-up research using datasets with respiratory information would significantly improve interpretation regarding the contribution of nuisance variables to the frequency spectra. Further, the authors would suggest that any researcher utilizing multi-band for data collection include methods to record this respiratory information.

Chen et al. 2017 identified a relationship in which stepwise nuisance regression may negatively influence BOLD signal at higher frequencies. The authors attempted to avoid this problem by applying the filtering directly to the data with no nuisance regressions, but included covariates and nuisance variables in the final model. Variance and effect sizes for motion and the motion by bin interactions were relatively small, with a minimal number of the LMEs returning as statistically significant. However, it should be noted that partial 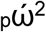 does not take correlations for within-participant measures into account, producing 25-50% inflation within the total 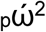. While generalized 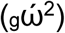 addresses this limitation, its computation is not intuitive for models with many factors. While the authors acknowledge this limitation, it is worth noting that even with a conservative 50-75% reduction in 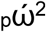, the effect size of frequency bin is still quite large 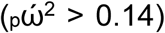, and the nuisance variable 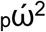 values are small. As such, the authors conclude that while motion may have influenced the signal within some connections, the effect seems minimal within the overall results.

It is worth noting that the FNC patterns and *L* increasing up to the 0.31-0.46 Hz frequency bin, then decreasing as the frequency range climbas likely varies depending on the type (single vs. multi-band) and TR of the data. As such, while the authors conclude that visually randomness and FNC patterns appear to be modular even with the increase and decrease relative to the 0.31-0.46 Hz bin, generalization of this pattern across variable TRs and acquisition parameters have yet to be established.

The method used to remove frequency information outside of the “bins” of interest is not a true digital filter of the data, but rather a regression model where all signals outside of the bin of interest are treated as nuisance information. While this approach is advocated by research groups performing nuisance regression with fMRI data (Chen et al., 2017; Lindquist et al., 2019), the effect on the frequency from a phase perspective, the degree of ripple, and the speed of the transition from passband to stopband have not been studied in detail. The effects of the nuisance-regression approach will need to be examined before broader conclusions are made.

Even when considering these limitations, the results of this research indicate variability in higher-frequency components that have the potential to be developmentally, cognitively, and diagnostically useful. When spatially constraining ICAs were derived from full-spectrum data to selective “bins” across higher frequencies, large differences in the FNC measures and randomness were identified even when the effects of other variables are regressed out. Such an approach would be useful in the identification of both neuronal signal of interest, as well as sources of non-neuronal artifact in future studies. Identifying such frequency-specific components has the potential to improve heuristic classification approaches, and the authors encourage the development of data-driven measures for BOLD filter design and applications with this goal in mind.

## Supporting information

Supplementary Information: Model Results

Reference Components and Custom Templates

## Author contributions

**T. DeRamus:** Performed ICA and regression analyses, prepared manuscript, constructed figures, adapted workflow to work with *SLURM* scheduler.

**Ashkan Faghiri:** Provided significant input regarding signal processing, aided in workflow development for filtering/nuisance regression.

**Armin Iraji:** Significantly contributed to manuscript proofing and implementation of ICA, one of three authors whom visually classified components by domain.

**Oktay Agcaoglu:** Performed initial analysis on full-spectrum dataset which generated the MOO-ICAR priors. One of the three authors who visually classified component domains.

**Victor M. Vergara:** Developed the method and *MATLAB* workflow for the randomness *L* calculation.

**Rogers Silva:** Aided in the development and implementation of the experimental steps and workflow necessary to assess the hypotheses of interest. Provided conceptual and pratical explanations for MOO-ICA implementation.

**Zening Fu:** Developed the original *MATLAB* workflow used for the MOO-ICA and FNC matrices. One of the three collaborators who performed visual classification of components by functional domain.

**Harshvardhan Gazula:** Provided valuable feedback regarding workflow construction, implementation, and interpretation.

**Julia M. Stephen, Tony Wilson, Yu-Ping Wang, Vince Calhoun:** Principal investigators in the collaborative developmental chronnecto-genomics (Dev-CoG) initiative. Collected and provided the imaging and demographic data. The NSF grant awarded to Dev-CoG collaborators and the NIBIB and NIMH research grants funded the analysis presented in this manuscript.

## Funding

National Science Foundation: #1539067

National Institute of General Medical Sciences: P20GM103472, P20GM130447

National Institute of Biomedical Imaging and Bioengineering: R01EB020407

National Institute of Mental Health: R01MH121101

## Acknowledgements

The authors would like to acknowledge the high-performance computing staff at Georgia State University for their implementation and maintenance of all resources used to conduct this study, the open-source tools and communities that facilitated these analyses *(R* core team and package developers, Python and Seaborn developers, *SLURM* scheduler, *AFNI, ANTs*, and *FSL* teams), Eswar Damaraju for image preprocessing support, and Srinivas Rachakonda for his continued development and maintenance of the *GIFT* toolbox.

## References

Agcaoglu, O., Wilson, T. W., Wang, Y.-P., Stephen, J., & Calhoun, V. D. (2019). Resting state connectivity differences in eyes open versus eyes closed conditions. Human Brain Mapping, 40(8), 2488–2498. doi: 10.1002/hbm.24539

Avants, B B, Epstein, C. L., Grossman, M., & Gee, J. C. (2008). Symmetric diffeomorphic image registration with cross-correlation: evaluating automated labeling of elderly and neurodegenerative brain. Medical Image Analysis, 12(1), 26–41. doi: 10.1016/j.media.2007.06.004

Avants, Brian B, Tustison, N. J., Song, G., Cook, P. A., Klein, A., & Gee, J. C. (2011). A reproducible evaluation of ANTs similarity metric performance in brain image registration. Neuroimage, 54(3), 2033–2044. doi: 10.1016/j.neuroimage.2010.09.025

Avants, Brian B, Tustison, N. J., Wu, J., Cook, P. A., & Gee, J. C. (2011). An open source multivariate framework for n-tissue segmentation with evaluation on public data. Neuroinformatics, 9(4), 381–400. doi: 10.1007/s12021-011-9109-y

Avants, Brian B., Tustison, N., & Song, G. (2009). Advanced Normalization Tools (ANTS). Insight Journal, 1–35.

Bates, D., Mächler, M., Bolker, B., & Walker, S. (2015). Fitting linear mixed-effects models using lme4. Journal of statistical software, 67(1), 1–48. doi: 10.18637/jss.v067.i01

Bell, A. J., & Sejnowski, T. J. (1995). An information-maximization approach to blind separation and blind deconvolution. Neural Computation, 7(6), 1129–1159. doi: 10.1162/neco.1995.7.6.1129

Biswal, B., DeYoe, E. A., & Hyde, J. S. (1996). Reduction of physiological fluctuations in fMRI using digital filters. Magnetic Resonance in Medicine, 35(1), 107–113. doi: 10.1002/mrm.1910350114

Burger, H. C., & Schuler, C. J. (2012). Image denoising: Can plain neural networks compete with BM3D? 2012 IEEE conference on ….

Burgess, G. C., Kandala, S., Nolan, D., Laumann, T. O., Power, J. D., Adeyemo, B., … Barch, D. M. (2016). Evaluation of Denoising Strategies to Address Motion-Correlated Artifacts in RestingState Functional Magnetic Resonance Imaging Data from the Human Connectome Project. Brain connectivity, 6(9), 669–680. doi: 10.1089/brain.2016.0435

Caballero-Gaudes, C., & Reynolds, R. C. (2017). Methods for cleaning the BOLD fMRI signal. Neuroimage, 154, 128–149. doi: 10.1016/j.neuroimage.2016.12.018

Calhoun, V. D., Adali, T., Pearlson, G. D., & Pekar, J. J. (2001). A method for making group inferences from functional MRI data using independent component analysis. Human Brain Mapping, 14(3), 140–151. doi: 10.1002/hbm.1048

Chen, J. E., & Glover, G. H. (2015). BOLD fractional contribution to resting-state functional connectivity above 0.1 Hz. Neuroimage, 107, 207–218. doi: 10.1016/j.neuroimage.2014.12.012

Chen, J. E., Jahanian, H., & Glover, G. H. (2017). Nuisance Regression of High-Frequency Functional Magnetic Resonance Imaging Data: Denoising Can Be Noisy. Brain connectivity, 7(1), 13–24. doi: 10.1089/brain.2016.0441

Cordes, D., Haughton, V. M., Arfanakis, K., Carew, J. D., Turski, P. A., Moritz, C. H., … Meyerand, M. E. (2001). Frequencies contributing to functional connectivity in the cerebral cortex in “restingstate” data. American Journal of Neuroradiology, 22(7), 1326–1333.

Cox, R. W. (1996). AFNI: software for analysis and visualization of functional magnetic resonance neuroimages. Computers and biomedical research, an international journal, 29(3), 162–173. doi: 10.1006/cbmr.1996.0014

Du, Y., & Fan, Y. (2013). Group information guided ICA for fMRI data analysis. Neuroimage, 69, 157–197. doi: 10.1016/j.neuroimage.2012.11.008

Erhardt, E. B., Rachakonda, S., Bedrick, E. J., Allen, E. A., Adali, T., & Calhoun, V. D. (2011). Comparison of multi-subject ICA methods for analysis of fMRI data. Human Brain Mapping, 32(12), 2075–2095. doi: 10.1002/hbm.21170

Fair, D. A., Miranda-Dominguez, O., Snyder, A. Z., Perrone, A., Earl, E. A., Van, A. N., … Dosenbach, N. U. F. (2020). Correction of respiratory artifacts in MRI head motion estimates. Neuroimage, 208, 116400. doi: 10.1016/j.neuroimage.2019.116400

Ferrarini, L., Veer, I. M., Baerends, E., van Tol, M.-J., Renken, R. J., van der Wee, N. J. A., … Milles, J. (2009). Hierarchical functional modularity in the resting-state human brain. Human Brain Mapping, 30(7), 2220–2231. doi: 10.1002/hbm.20663

Friston, K. J., Holmes, A. P., Worsley, K. J., Poline, J. P., Frith, C. D, & Frackowiak, R. S. J. (1994). Statistical parametric maps in functional imaging: A general linear approach. Human Brain Mapping, 2(4), 189–210. doi: 10.1002/hbm.460020402

Girko, V. L. (1985). Circular Law. Theory of Probability & Its Applications, 29(4), 694–706. doi: 10.1137/1129095

Gratton, C., Coalson, R. S., Dworetsky, A., Adeyemo, B., Laumann, T. O., Wig, G. S., … Campbell, M. C. (2019). Removal of high frequency contamination from motion estimates in single-band fMRI saves data without biasing functional connectivity. BioRxiv. doi: 10.1101/837161

Himberg, J., & Hyvarinen, A. (2003). Icasso: software for investigating the reliability of ICA estimates by clustering and visualization. In 2003 IEEE XIII Workshop on Neural Networks for Signal Processing (IEEE Cat. No.03TH8718) (pp. 259–268). IEEE. doi: 10.1109/NNSP.2003.1318025

Holm, S. (1979). A simple sequentially rejective multiple test procedure. Scandinavian journal of statistics, 65–70.

Jaeger, B. (2017). r2glmm: Computes R Squared for Mixed (Multilevel) Models.

Jenkinson, M., Beckmann, C. F., Behrens, T. E. J., Woolrich, M. W., & Smith, S. M. (2012). FSL. Neuroimage, 62(2), 782–790. doi: 10.1016/j.neuroimage.2011.09.015

Jo, H. J., Gotts, S. J., Reynolds, R. C., Bandettini, P. A., Martin, A., Cox, R. W., & Saad, Z. S. (2013). Effective Preprocessing Procedures Virtually Eliminate Distance-Dependent Motion Artifacts in Resting State FMRI. Journal of applied mathematics, 2013. doi: 10.1155/2013/935154

Johnson, P. C. (2014). Extension of Nakagawa & Schielzeth’s R2GLMM to random slopes models. Methods in ecology and evolution / British Ecological Society, 5(9), 944–946. doi: 10.1111/2041-210X.12225

Klein, A., Andersson, J., Ardekani, B. A., Ashburner, J., Avants, B., Chiang, M.-C., … Parsey, R. V. (2009). Evaluation of 14 nonlinear deformation algorithms applied to human brain MRI registration. Neuroimage, 46(3), 786–802. doi: 10.1016/j.neuroimage.2008.12.037

Kuznetsova, A., Brockhoff, P. B., & Christensen, R. H. B. (2017). lmertest package: tests in linear mixed effects models. Journal of statistical software, 82(13). doi: 10.18637/jss.v082.i13

Lenth, R. (2019). emmeans: Estimated Marginal Means, aka Least-Squares Means.

Li, X.-L., & Adali, T. (2010a). Blind spatiotemporal separation of second and/or higher-order correlated sources by entropy rate minimization. In 2010 IEEE International Conference on Acoustics, Speech and Signal Processing (pp. 1934–1937). IEEE. doi: 10.1109/ICASSP.2010.5495311

Li, X.-L., & Adali, T. (2010b). Independent component analysis by entropy bound minimization. IEEE Transactions on Signal Processing, 58(10), 5151–5164. doi: 10.1109/TSP.2010.2055859

Liang, B., Zhang, D., Wen, X., Xu, P., Peng, X., Huang, X., … Huang, R. (2014). Brain spontaneous fluctuations in sensorimotor regions were directly related to eyes open and eyes closed: evidences from a machine learning approach. Frontiers in Human Neuroscience, 8, 645. doi: 10.3389/fnhum.2014.00645

Lindquist, M. A., Geuter, S., Wager, T. D., & Caffo, B. S. (2019). Modular preprocessing pipelines can reintroduce artifacts into fMRI data. Human Brain Mapping, 40(8), 2358–2376. doi: 10.1002/hbm.24528

Liu, D., Dong, Z., Zuo, X., Wang, J., & Zang, Y. (2013). Eyes-open/eyes-closed dataset sharing for reproducibility evaluation of resting state fMRI data analysis methods. Neuroinformatics, 11(4), 469–476. doi: 10.1007/s12021-013-9187-0

Long, J. A. (2019). interactions: Comprehensive, User-Friendly Toolkit for Probing Interactions.

Luo, D., Ganesh, S., & Koolaard, J. (2020). predictmeans: Calculate Predicted Means for Linear Models.

Ma, S., Correa, N. M., Li, X.-L., Eichele, T., Calhoun, V. D., & Adali, T. (2011). Automatic identification of functional clusters in FMRI data using spatial dependence. IEEE Transactions on Bio-Medical Engineering, 58(12), 3406–3417. doi: 10.1109/TBME.2011.2167149

Marcenko, V. A., & Pastur, L. A. (1967). Distribution of eigenvalues for some sets of random matrices. Mathematics of the USSR-Sbornik, 1(4), 457–483. doi: 10.1070/SM1967v001n04ABEH001994

Morgan, V. L., Rogers, B. P., & Abou-Khalil, B. (2015). Segmentation of the thalamus based on BOLD frequencies affected in temporal lobe epilepsy. Epilepsia, 56(11), 1819–1827. doi: 10.1111/epi.13186

Nakagawa, S., & Schielzeth, H. (2013). A general and simple method for obtaining R2 from generalized linear mixed-effects models.

Newman, M. E. J. (2006). Modularity and community structure in networks. Proceedings of the National Academy of Sciences of the United States of America, 103(23), 8577–8582. doi: 10.1073/pnas.0601602103

Niazy, R. K., Xie, J., Miller, K., Beckmann, C. F., & Smith, S. M. (2011). Spectral characteristics of resting state networks. Progress in Brain Research, 193, 259–276. doi: 10.1016/B978-0-444-53839-0.00017-X

Patriat, R., Molloy, E. K, Meier, T. B., Kirk, G. R., Nair, V. A., Meyerand, M. E., … Birn, R. M. (2013). The effect of resting condition on resting-state fMRI reliability and consistency: a comparison between resting with eyes open, closed, and fixated. Neuroimage, 78, 463–473. doi: 10.1016/j.neuroimage.2013.04.013

R Core team. (2015). R Core Team. R: A Language and Environment for Statistical Computing. R Foundation for Statistical Computing, Vienna, Austria. ISBN 3-900051-07-0, URL http://www.R-project.org/., 55, 275-286.

Rubinov, M., & Sporns, O. (2010). Complex network measures of brain connectivity: uses and interpretations. Neuroimage, 52(3), 1059–1069. doi: 10.1016/j.neuroimage.2009.10.003

Sanchez, C. E., Richards, J. E., & Almli, C. R. (2012). Neurodevelopmental MRI brain templates for children from 2 weeks to 4 years of age. Developmental Psychobiology, 54(1), 77–91. doi: 10.1002/dev.20579

Satterthwaite, T. D., Elliott, M. A., Gerraty, R. T., Ruparel, K., Loughead, J., Calkins, M. E., .. Wolf, D. H. (2013). An improved framework for confound regression and filtering for control of motion artifact in the preprocessing of resting-state functional connectivity data. Neuroimage, 64, 240–256. doi: 10.1016/j.neuroimage.2012.08.052

Smith, S. M., Jenkinson, M., Woolrich, M. W., Beckmann, C. F., Behrens, T. E. J., Johansen-Berg, H.,. Matthews, P. M. (2004). Advances in functional and structural MR image analysis and implementation as FSL. Neuroimage, 23 Suppl 1, S208–19. doi: 10.1016/j.neuroimage.2004.07.051

Sours, C., Chen, H., Roys, S., Zhuo, J., Varshney, A., & Gullapalli, R. P. (2015). Investigation of Multiple Frequency Ranges Using Discrete Wavelet Decomposition of Resting-State Functional Connectivity in Mild Traumatic Brain Injury Patients. Brain connectivity, 5(7), 442–450. doi: 10.1089/brain.2014.0333

Sporns, O. (2003). Graph Theory Methods for the Analysis of Neural Connectivity Patterns. In Neuroscience Databases (pp. 171–185). Boston, MA: Springer US. doi: 10.1007/978-1-4615-1079-6_12

Team, R. C. (2019). R: A Language and Environment for Statistical Computing.

Torchiano, M. (2016). Effsize - A Package For Efficient Effect Size Computation. Zenodo. doi: 10.5281/zenodo.1480624

Trotter, H. F. (1984). Eigenvalue distributions of large Hermitian matrices; Wigner’s semi-circle law and a theorem of Kac, Murdock, and Szegö. Advances in mathematics, 54(1), 67-82. doi: 10.1016/0001-8708(84)90037-9

Vergara, V. M., & Calhoun, V. (2016). Randomness in resting state functional connectivity matrices. Conference Proceedings: Annual International Conference of the IEEE Engineering in Medicine and Biology Society, 2016, 5563–5566. doi: 10.1109/EMBC.2016.7591987

Vergara, V. M., & Calhoun, V. D. (2020). Filtered correlation and allowed frequency spectra in dynamic functional connectivity. Journal of Neuroscience Methods, 108837. doi: 10.1016/j.jneumeth.2020.108837

Vergara, V. M., Yu, Q., & Calhoun, V. D. (2018a). A method to assess randomness of functional connectivity matrices. Journal of Neuroscience Methods, 303, 146–158. doi: 10.1016/j.jneumeth.2018.03.015

Vergara, V. M., Yu, Q., & Calhoun, V. D. (2018b). Graph modularity and randomness measures: A comparative study. In 2018 IEEE Southwest Symposium on Image Analysis and Interpretation (SSIAI) (pp. 33–36). IEEE. doi: 10.1109/SSIAI.2018.8470322

Wang, J., Zhang, Z., Ji, G.-J., Xu, Q., Huang, Y., Wang, Z., … Lu, G. (2015). Frequency-Specific Alterations of Local Synchronization in Idiopathic Generalized Epilepsy. Medicine, 94(32), e1374. doi: 10.1097/MD.0000000000001374

Wei, J., Chen, T., Li, C., Liu, G., Qiu, J., & Wei, D. (2018). Eyes-Open and Eyes-Closed Resting States With Opposite Brain Activity in Sensorimotor and Occipital Regions: Multidimensional Evidences From Machine Learning Perspective. Frontiers in Human Neuroscience, 12, 422. doi: 10.3389/fnhum.2018.00422

Wilcoxon, F. (1946). Individual comparisons of grouped data by ranking methods. Journal of Economic Entomology, 39, 269.

Yoo, A. B., Jette, M. A., & Grondona, M. (2003). SLURM: simple linux utility for resource management. In D. Feitelson, L. Rudolph, & U. Schwiegelshohn (eds.), Job scheduling strategies for parallel processing (Vol. 2862, pp. 44-60). Berlin, Heidelberg: Springer Berlin Heidelberg. doi: 10.1007/10968987_3

Yu, Q., Plis, S. M., Erhardt, E. B., Allen, E. A., Sui, J., Kiehl, K. A., … Calhoun, V. D. (2011). Modular Organization of Functional Network Connectivity in Healthy Controls and Patients with Schizophrenia during the Resting State. Frontiers in Systems Neuroscience, 5, 103. doi: 10.3389/fnsys.2011.00103

Yuan, B.-K., Wang, J., Zang, Y.-F., & Liu, D.-Q. (2014). Amplitude differences in high-frequency fMRI signals between eyes open and eyes closed resting states. Frontiers in Human Neuroscience, 8, 503. doi: 10.3389/fnhum.2014.00503

Zou, Q., Long, X., Zuo, X., Yan, C., Zhu, C., Yang, Y., … Zang, Y. (2009). Functional connectivity between the thalamus and visual cortex under eyes closed and eyes open conditions: a resting-state fMRI study. Human Brain Mapping, 30(9), 3066–3078. doi: 10.1002/hbm.20728

Zou, Q., Yuan, B.-K., Gu, H., Liu, D., Wang, D. J. J., Gao, J.-H.,. Zang, Y.-F. (2015). Detecting static and dynamic differences between eyes-closed and eyes-open resting states using ASL and BOLD fMRI. Plos One, 10(3), e0121757. doi: 10.1371/journal.pone.0121757

