## Supplementary figures and images for "Modular and state-relevant connectivity in high-frequency resting-state BOLD fMRI data: An independent component analysis"

### icatb_network_summary_01.png

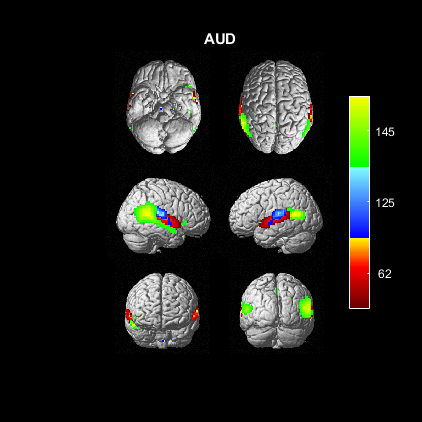

### icatb_network_summary_02.png

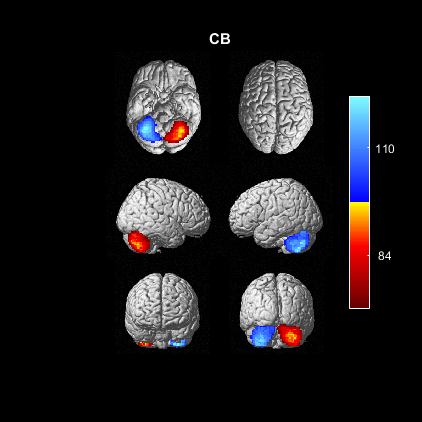

### icatb_network_summary_03.png

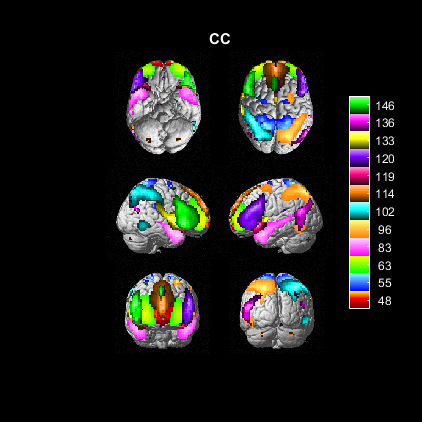

### icatb_network_summary_04.png

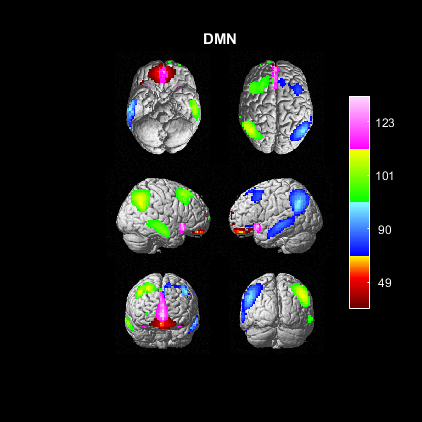

### icatb_network_summary_05.png

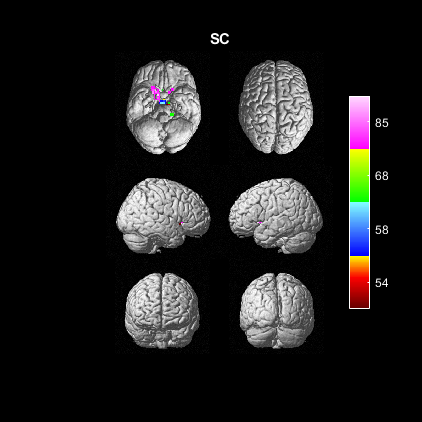

### icatb_network_summary_06.png

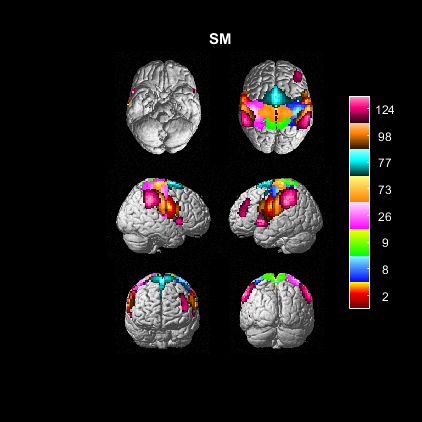

### icatb_network_summary_07.png

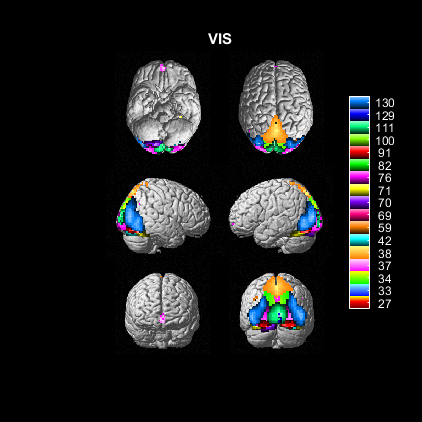

### icatb_network_summary_08.png

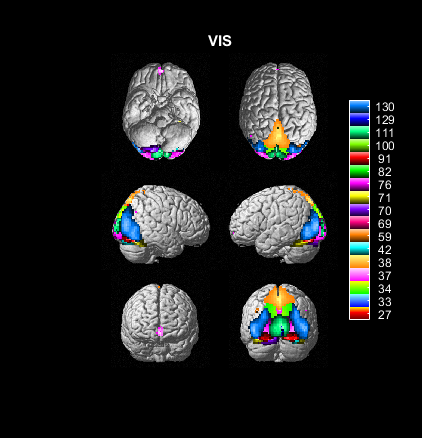

### icatb_network_summary_09.png

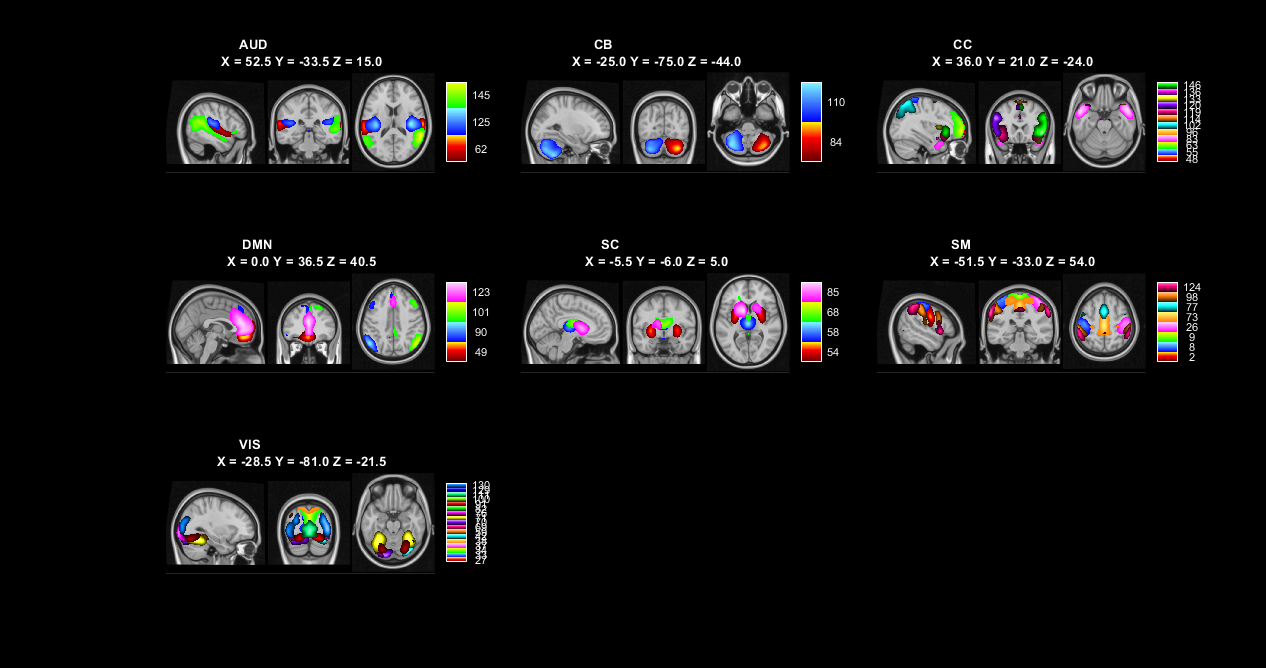

### icatb_network_summary_10.png

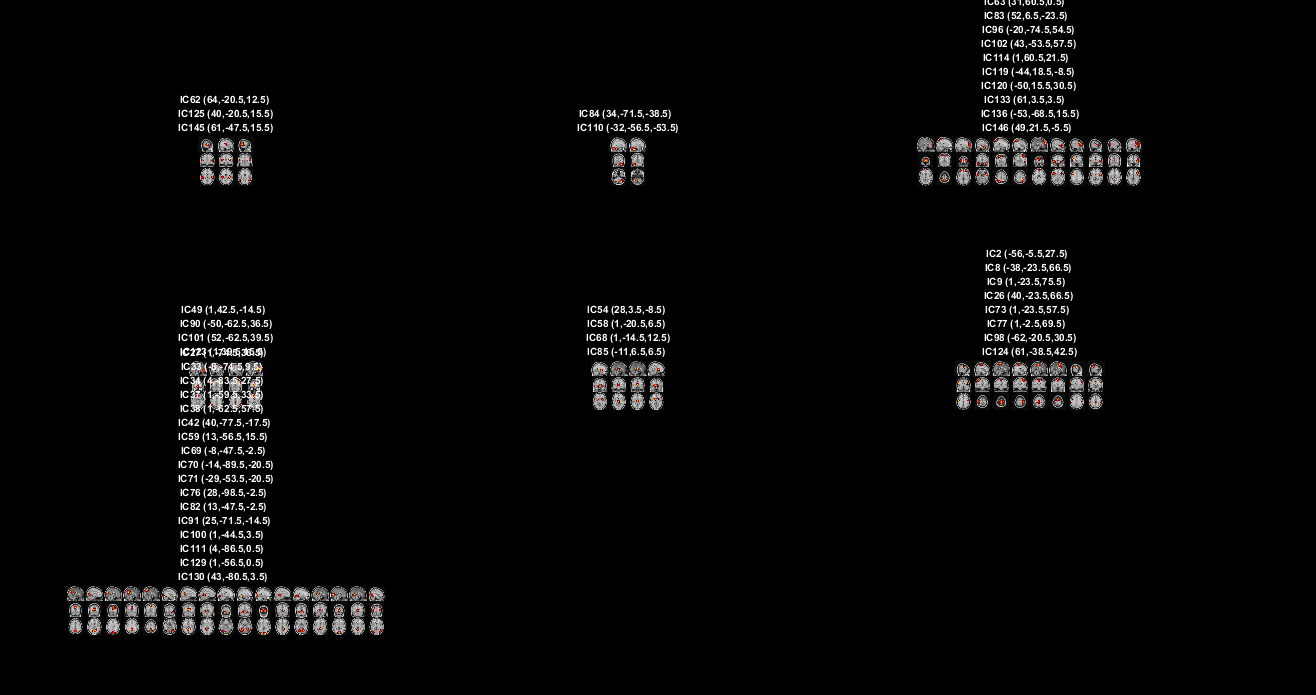

### Supplemental Figure 1

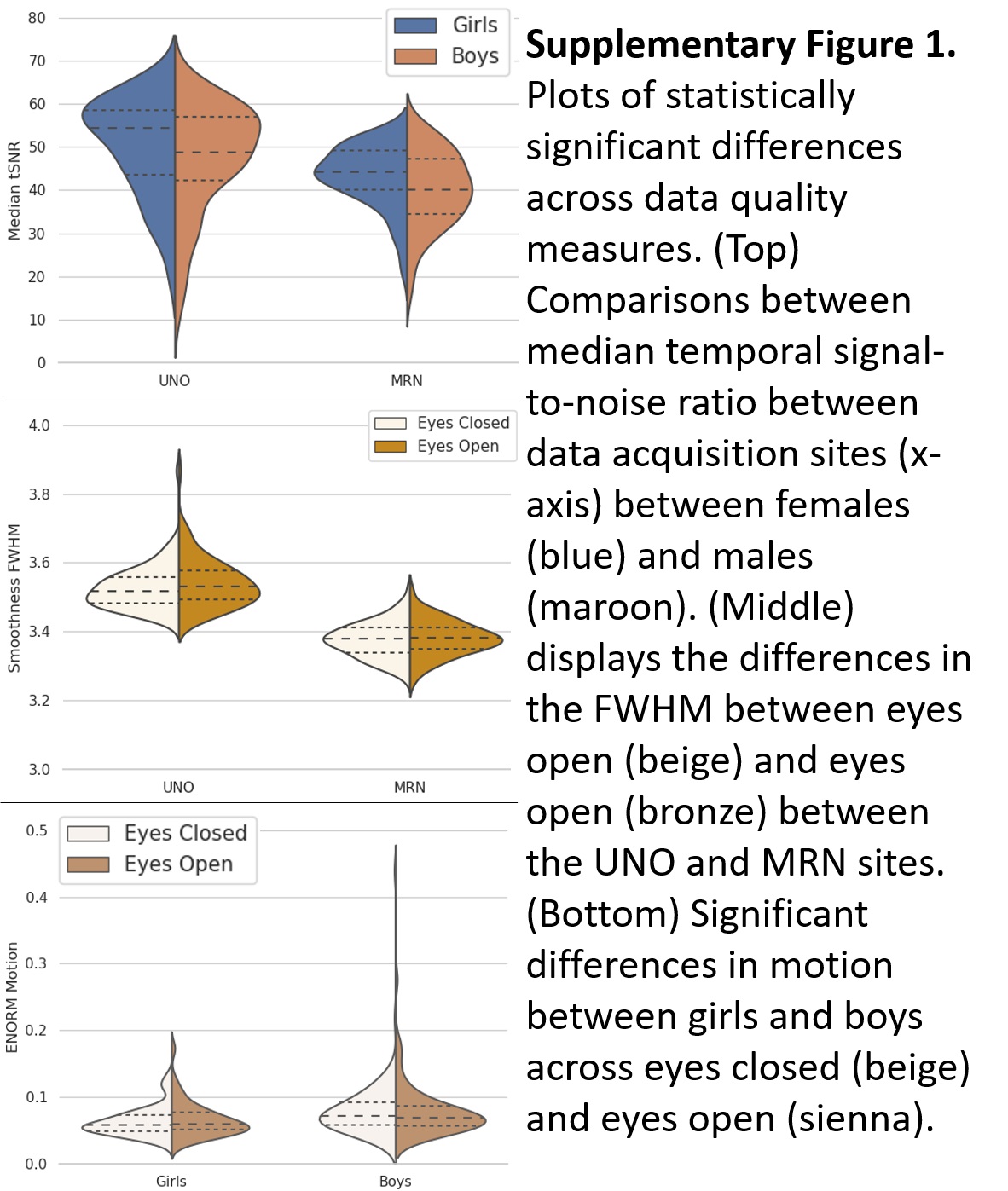

### Supplementary Figure 2

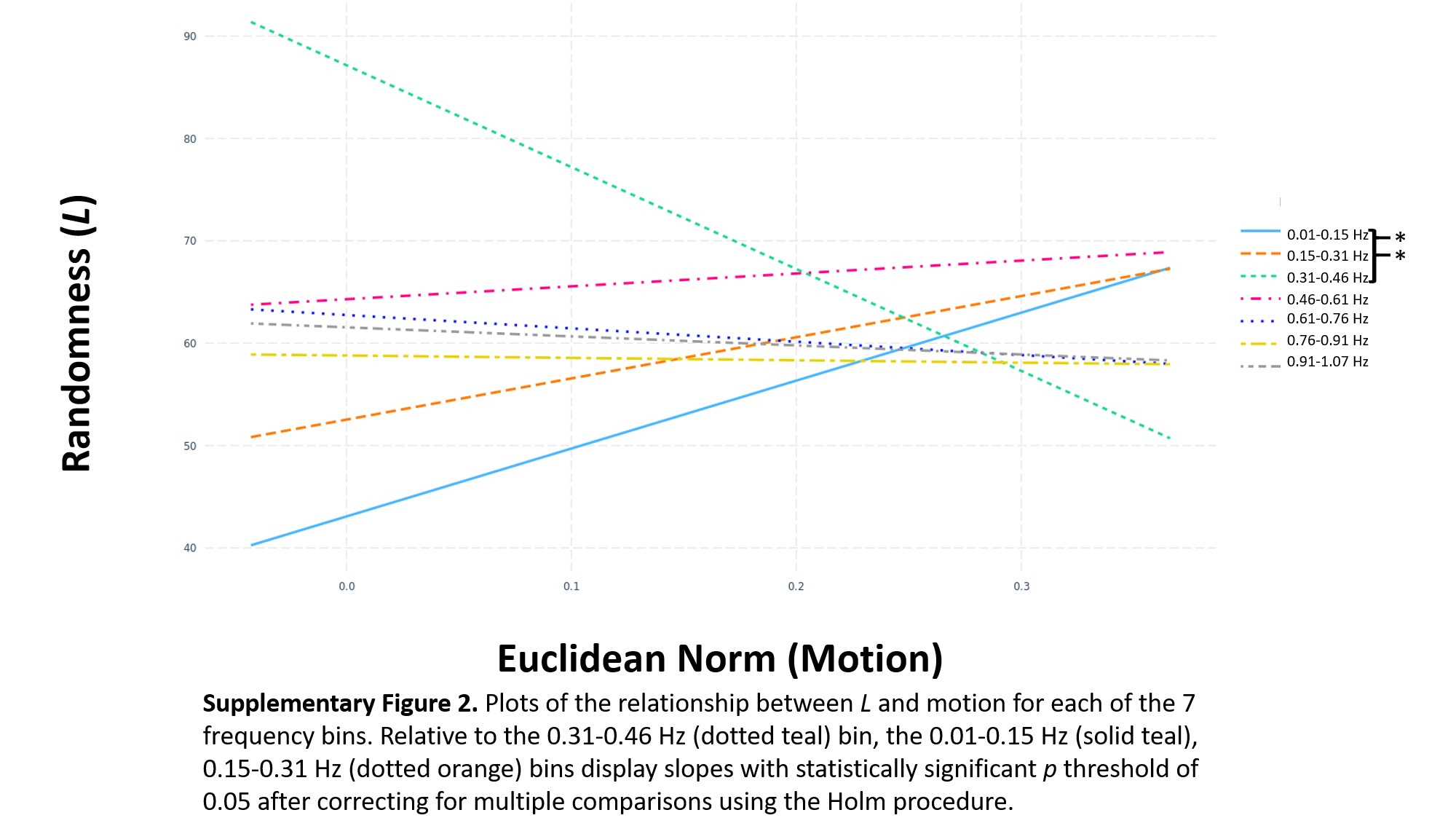
