## Supplementary material for "Modular and state-relevant connectivity in high-frequency resting-state BOLD fMRI data: An independent component analysis": Reference Components and Custom Templates: icatb_network_summary.html

### Contents

- Rendering: Multiple components are rendered on surface of a 'standard' brain
- FNC correlations: Correlations are visualized in a matrix plot
- Connectogram view - FNC correlations are shown using bezier curves and thumbnails of spatial maps are shown in a circle. Components within the same network are shown in the same color.
- Multiple components are displayed in a composite plot. Orthogonal slices are shown.
- Stacked ortho slices are shown for each component in the network. Title shows component numbers plotted from top to bottom

### Rendering: Multiple components are rendered on surface of a 'standard' brain

      

### FNC correlations: Correlations are visualized in a matrix plot

### Connectogram view - FNC correlations are shown using bezier curves and thumbnails of spatial maps are shown in a circle. Components within the same network are shown in the same color.

### Multiple components are displayed in a composite plot. Orthogonal slices are shown.

 

### Stacked ortho slices are shown for each component in the network. Title shows component numbers plotted from top to bottom

Published with MATLAB® R2019a
